# Axial Rotation Comprises Concurrent Twisting and Bending, Each Differentially Regulated by TGF-β Signaling

**DOI:** 10.1101/2025.10.23.683873

**Authors:** Yuki S. Kogure, Satoru Okuda, Kotaro Oka, Kohji Hotta

## Abstract

Axial rotation (AR), a morphogenetic movement that reshapes the body axis, is widely observed in chordates, including mice and rats. AR involves complex three-dimensional deformations; however, its geometric characteristics and regulatory mechanisms remain poorly understood. Here, using the chordate *Ciona robusta* (*Ciona intestinalis* type A), we demonstrate that AR consists of two components—leftward bending and clockwise twisting along the anterior–posterior axis—differentially regulated by TGF-β signaling. A comparison between chorionated and dechorionated embryos revealed that dechorionation randomized the bending direction, while twisting remained consistently clockwise. Inhibition of TGF-β signaling with SB431542 randomized both deformations. Quantitative analysis of twisting angles indicated uniform clockwise twisting along the axis, peaking during the tailbud stage and proceeding in the tail region, independent of the tip, trunk, or myofibril patterning. Although overall twisting was reduced under TGF-β inhibition, the tail exhibited disorganized twisting. The sum of absolute twisting-angle differences in every 10 μm remained comparable to the wild type (WT). This suggests that twisting is intrinsically generated, while TGF-β signaling aligns local twisting into a coordinated global direction. Our findings dissected the mechanisms of AR in *Ciona* and highlight the multilayered regulation underlying the morphogenesis of the chordate body plan and providing a foundation for understanding its biomechanical and molecular bases.

## 1. Introduction

During organogenesis, chordate embryos exhibit similar morphologies and highly conserved gene expression profiles (Haeckel, 1874; Irie and Kuratani, 2011). A notable morphological transition at this stage is axial rotation (AR), which causes a significant change in body posture. In mice and rats, embryos first display dorsiflexion (bending toward the dorsal side) and subsequently undergo ventroflexion (bending toward the ventral side) (Fujinaga et al., 1995; Kaufman, 1992; Poelmann et al., 1987). AR has also been observed in chicks and in reptiles such as alligators and snakes (Ferguson, 1981; Hamburger and Hamilton, 1951; Poelmann et al., 1987; Zehr, 1962). The presence of AR across multiple vertebrate lineages suggests that it may represent an evolutionarily conserved developmental feature. The significance of AR is underscored by the lethality observed in many AR-defective mutants (Chatterjee et al., 2007; Constam and Robertson, 2000; Faisst et al., 2002; Imamoto and Soriano, 1993; Nada et al., 1993; Roebroek et al., 1998).

AR has primarily been studied using mice (and rats) and chicks as model organisms. In mouse and rat embryos, AR has been described in detail by (Kaufman, 1992) and (Fujinaga et al., 1995). Kaufman proposed a simplified model of mouse embryonic development, which was (Fujinaga et al., 1995). According to this model, “the upper and lower bodies remain parallel as the lower body makes a horizontal rotation around the dorsoventral axis of the upper body” ((Fujinaga et al., 1995); Suppl. Fig. 1). In rat embryos, AR is subdivided into three movements: twisting of the upper body (stage 12/s7–8), the middle body (stage 13/s11–12), and the lower body (stage 14/s15–16) (Fujinaga et al., 1995). In chick embryos, AR initiates with anterior head turning to the left and then progresses caudally (Hamburger and Hamilton, 1951; Poelmann et al., 1987).

Based on these detailed morphological observations, AR can be considered a form of left-right symmetry breaking. Nodal signaling is a key pathway regulating such left-right asymmetries and is well known for establishing visceral asymmetries, such as heart looping and stomach situs (Levin et al., 1997, 1995; Lowe et al., 2001, 1996). In mice and chicks, asymmetric Nodal expression correlates with AR (Collignon et al., 1996; Levin et al., 1997; Logan et al., 1998). However, studies in chicks suggest that AR is independent of visceral laterality (Levin et al., 1997). These observations highlight the need to analyze AR and visceral asymmetries as separate processes. Because AR and organ asymmetries arise concurrently, analyzing AR independently has been challenging. AR-defective mutants often exhibit additional morphological defects (Chatterjee et al., 2007; Constam and Robertson, 2000; Faisst et al., 2002; Imamoto and Soriano, 1993; Nada et al., 1993; Roebroek et al., 1998), complicating the interpretation of AR-specific defects. Despite its evolutionary and developmental significance, the mechanisms underlying AR remain largely unknown. Moreover, most previous studies have relied on qualitative descriptions, underscoring the need for quantitative analyses.

To address these issues, this study focused on the ascidian *Ciona robusta* (also known as *Ciona intestinalis* type A). During the tailbud stage, embryos reverse tail bending from the ventral to the dorsal side (Hotta et al., 2007) and the tail epidermis is organized into eight rows of epidermal cells (Hotta et al., 2007; Kogure et al., 2022). Left–right asymmetric tail bending occurs at stage (st.) 23 (Palmquist and Davidson, 2017). Our detailed observations of embryo morphology and epidermal row alignment across multiple time points revealed posture transitions resembling AR in rodents (Fujinaga et al., 1995; Kaufman, 1992). At these stages, major visceral organs have not yet formed (Hotta et al., 2007; Nakamura et al., 2012), allowing AR to be examined independently. Furthermore, due to their small cell number and simple body plan (Nakamura et al., 2012), *Ciona* tailbud embryos serve as an ideal model for studying AR. Here, we demonstrate that AR consists of two distinct components: bending and twisting. While bending direction is regulated by the Nodal expression pattern (Nishide et al., 2012; Palmquist and Davidson, 2017), we show that twisting direction appears independent of this pattern. We also show that twisting direction depends on transforming growth factor β (TGF-β) signaling. To dissect these components, we established a method to quantify twisting independently of bending, revealing that TGF-β signaling aligns local twisting into a coherent global pattern. This study provides a valuable foundation for elucidating the biomechanical and molecular bases of AR.

## 2. Materials and Methods

### 2.1 Ascidian Samples

Adult *Ciona* were obtained from the Maizuru Fisheries Research Station (Kyoto University), Onagawa Field Center (Tohoku University), and Misaki Marine Biological Station (University of Tokyo) through the National Bio-Resource Project (Japan). Eggs were collected by dissecting gonoducts, then artificially dechorionated and fertilized as previously described (Hotta et al., 1999). For chorionated embryos, the chorion was manually removed after fixation. Fertilized eggs were incubated in filtered seawater at 16°C or 22°C until fixation in 4% paraformaldehyde (PFA) at 4°C overnight (Hotta et al., 2007). Samples were washed with phosphate-buffered saline (PBS) and stained overnight in PBS containing Alexa Fluor 546-conjugated phalloidin (1:50; Thermo Fisher). Developmental stages were assigned according to Hotta’s system (Hotta et al., 2020, 2007).

### 2.2 Inhibitor Treatment

Ascidian embryos were treated with 2.5 μM SB431542 (Nacalai Tesque) from the late gastrula or neurula stage until fixation.

To disrupt muscle fiber alignment (Ohtsuka and Okamura, 2007), embryos were cultured in seawater containing 1 mM d-tubocurarine chloride (Sigma-Aldrich) or 40 μM N-benzyl-p-toluenesulfonamide (BTS; CAS 1576-37-0; Tokyo Chemical Industry) from the gastrula stage until fixation, either at 16 hpf or at the swimming larva stage.

### 2.3 Quantification of Twisting

To independently evaluate tail twisting from overall body bending, we reconstructed three-dimensional (3D) images of phalloidin-stained embryos using confocal Z-stacks and developed an image analysis method. The reconstructed volume was segmented to extract a smooth central axis, which served as a reference for defining cross-sectional coordinate systems along the body axis. By resampling the original fluorescence data within these curvilinear coordinates, we generated “straightened” images that eliminated bending. These images were further transformed into two-dimensional (2D) surface and notochord projection maps, enabling clear visualization of epidermal rows and notochord cell boundaries. Finally, using the position of dorsal cells as a reference, we quantified twisting angles along the body axis, providing a robust measure of tissue torsion independent of embryo curvature. All image processing procedures were performed in MATLAB (MathWorks).

#### 2.3.1 Image acquisition and overlap alignment for large embryos

Z-stack images of phalloidin-stained embryos were acquired using Olympus FV1000 and FV4000 confocal laser scanning microscopes equipped with 40× objective lenses. When the embryo size exceeded the microscope’s field of view, images were captured in partially overlapping segments. Overlapping regions were identified by manually selecting slices with shared structures from each Z-stack and marking a representative pixel. The corresponding spatial coordinates (*x, y, z*) were calculated, and the segmented Z-stacks were merged into a continuous dataset encompassing the entire embryo.

#### 2.3.2 Segmentation and skeletonization

From the complete Z-stack, background fluorescence was removed by applying an intensity threshold that was manually adjusted to yield an accurate embryo silhouette. The image was then binarized to distinguish internal from external regions. A one-voxel-wide skeleton was obtained by thinning the binary mask while preserving its topology:

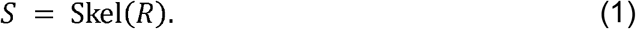

Where *R* ⊂ ℝ^3^ represents the segmented region, this skeleton provides a discrete approximation of the medial axis. The main branch of the skeleton was parameterized as a smooth centerline.

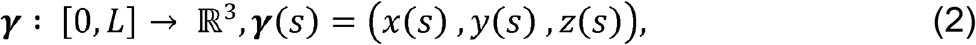

where *s* represents the arc length from the anterior to the posterior along the curve, and *L* is the total length.

#### 2.3.3 Local coordinate system and curvilinear transformation

At each point along the centerline, we considered the plane orthogonal to the tangent vector ***t***(*s*). This plane defines the cross-section used to analyze the local structure of the embryo. The unit tangent vector was defined as

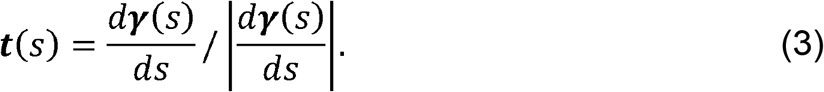

Within the plane orthogonal to ***t***(*s*), there remains freedom in choosing the orientation of the two in-plane basis vectors. To ensure a consistent orientation of these cross-sectional planes along the embryo, we defined the normal vector ***v***(*s*) to be as close as possible to the global *z*-axis ***e***_*z*_ = (0,0,1) :

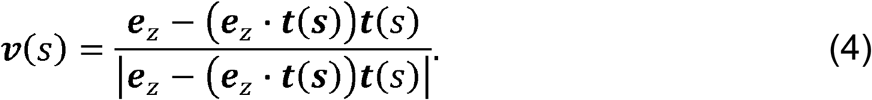

The remaining orthogonal direction ***u***(*s*)was then obtained as follows:

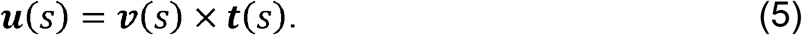

Thus, the triplet { ***t***(*s*), ***u***(*s*), ***v***(*s*)} forms an orthonormal basis along the curve, with ***v***(*s*) preferentially oriented toward the global *z*-axis. Using this local frame, the neighborhood of the centerline was described in cylindrical-like coordinates (*ρ, θ,s*), where *ρ* the radial distance from the centerline within the orthogonal plane, *θ* the angular direction in that plane, and *s* denotes the position along the centerline.

The original 3D image was represented as an intensity function *I* (*x,y,z*), where *l* denotes the fluorescence intensity at the standard 3D coordinates (*x,y,z*). This image was resampled into cylindrical-like coordinates to obtain a transformed image *J*(*ρ, θ,s*), which represents the fluorescence intensity in the straightened coordinate system.

#### 2.3.4 Surface unwrapping and notochord mapping (S-MAP and N-MAP)

To visualize the surface intensity distribution, we generated an unwrapped 2D map (S-MAP). In the curvilinear coordinate system (*ρ, θ,s*), the surface of the structure was identified as the boundary along the radial direction *ρ*, and the fluorescence intensity at this boundary was projected onto a plane with the *θ*- and *s*-axes. For the S-MAP, intensity values nearest to the surface of the structure along the *ρ* direction were extracted, and approximately 3 suface slices were combined using maximum intensity projection.

For the notochord projection image (N-MAP), intensity values nearest to the center of the structure along the *ρ* direction were extracted, and approximately 10 central slices were combined using maximum intensity projection. Thus, both S-MAP and N-MAP represent projection images defined in (*θ,s*) coordinates, providing a clear visualization of epidermal rows and notochord cell boundaries.

When rendering these maps, the *s*-axis was discretized according to the sequence of pixels along the centerline *S*, so that each pixel position on *S* corresponded to one step along the *s* -axis. In contrast, when calculating positions along the *s*-axis, the location was determined by summing the straight-line distances between consecutive points along the centerline in the original 3D space.

#### 2.3.5 Calculation of twisting angles

Notochord cells are aligned in a single chain along the anterior–posterior (AP) axis, with their radially oriented boundaries serving as discrete landmarks to determine the *s* -coordinate of epidermal cells. The position of the *i* -th boundary—between the *i* -th and (*i* +1)-th cells—is denoted as *s*_*i*_. On the N-MAP, notochord cell boundaries were first detected automatically and then corrected manually. —

The twisting angle was defined relative to the dorsal midline epidermal cells, which are also aligned in a chain-like manner along the *s*-axis. Two boundary lines form between these midline epidermal cells and their adjacent cells on the left and right sides. These boundaries were identified on the S-MAP using automatic detection, followed by manual correction.

To calculate the twisting angle, we smoothed the angular positions of each dorsal midline boundary within a defined range along the *s* -axis and established a reference point. We then averaged the values from the left and right boundaries. The averaging ranges and reference points were defined as follows:

i. Twisting angle at each notochord boundary (*φ* _*i*_): defined as *φ* _*i*_ =*θ* _*i*_ − *θ*_ref_ . Smoothing was performed over the interval *si*_−1_, ≤*s*< _*i*+1_. Here, =*θ* _*i*_ was defined as the smoothed angular position of the *i*th notochord boundary. The reference point was defined as the crossing point at the 5th notochord boundary (*s* = *s* _5_,) or a manually selected notochord boundary, and the smoothed angular position at this point was defined as *θ*_*ref*_.
ii. Twisting angles calculated every 10 μm (*ϕ* _*j*_): defined as *ϕ*_*j*_ =*θ* _*j*_− *θ*_ref_. Smoothing was performed over a 10 μm window along the *s*-axis. Here, *ϕ*_*j*_ was defined as the smoothed angular position at 10*j* μm from the reference point. The reference point was defined as the crossing point with the 5th notochord boundary (*s* = *s*_5_) before smoothing. The angular position at this point was defined as *θ*_ref_. When the 5th boundary was not identifiable, *θ*_ref_ was defined as the angular position at the *s* corresponding to the WT *s*_5_, which was manually selected based on the external morphology of the embryo.

To assess the extent of twisting independently of its direction, we defined the absolute twisting angle |Δ *ϕ*_*j*_ | as the absolute angular change in twisting between consecutive measurement points spaced at 10-µm intervals: |Δ *ϕ*_*j*_ | =| *ϕ* _*j* + 1_ − *ϕ* _*j*_ | . This metric quantifies the magnitude of local twisting regardless of whether it occurs clockwise or counterclockwise. The cumulative sum of absolute local twisting angles, *C*_*j*_, was then defined as

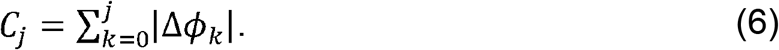

where *C*_*j*_ represents the accumulated twisting magnitude along the tail, starting from the reference point.

### 2.4 Surgical Excision of the Trunk and Tip

The trunk and tail tip were surgically removed at 13 hpf and incubated at 16ºC using glass needles (NARISHIGE, GC-1). The glass needles were fabricated with a P-97 glass needle puller (Sutter, California, USA) using the following settings: Heat = 574, P = 500, Pull = 100, Vel = 75, and Time = 150 (Matsumura et al., 2020).

## 3. Results

### 3.1 *Ciona robusta* Undergoes Axial Rotation During the Organogenesis Period

To investigate dynamic morphological changes during the tailbud stages, *Ciona* embryos were fixed at multiple developmental time points, and cell–cell boundaries were visualized by F-actin staining (Fig. 1A). Consistent with previous reports, embryos exhibited ventroflexion from st. 18 to st. 22 (Hotta et al., 2007; Kogure et al., 2022). From st. 23 to 25 (17 hpf), the embryos shifted their bending direction from the ventral to the lateral side (Fig. 1A; Suppl. Fig. 2A). This transition continued until at least st. 25 (19 hpf), with the bending direction shifting further to the dorsal side, resulting in dorsiflexion (Fig. 1A, st. 25; Suppl. Fig. 2A). In addition to earlier studies that focused on a single stage (Palmquist and Davidson, 2017), our observations across multiple time points suggest that left–right asymmetrical bending occurs as part of a continuous postural transition, representing an approximately 180° shift in bending direction from ventral to dorsal.

**Fig. 1.**
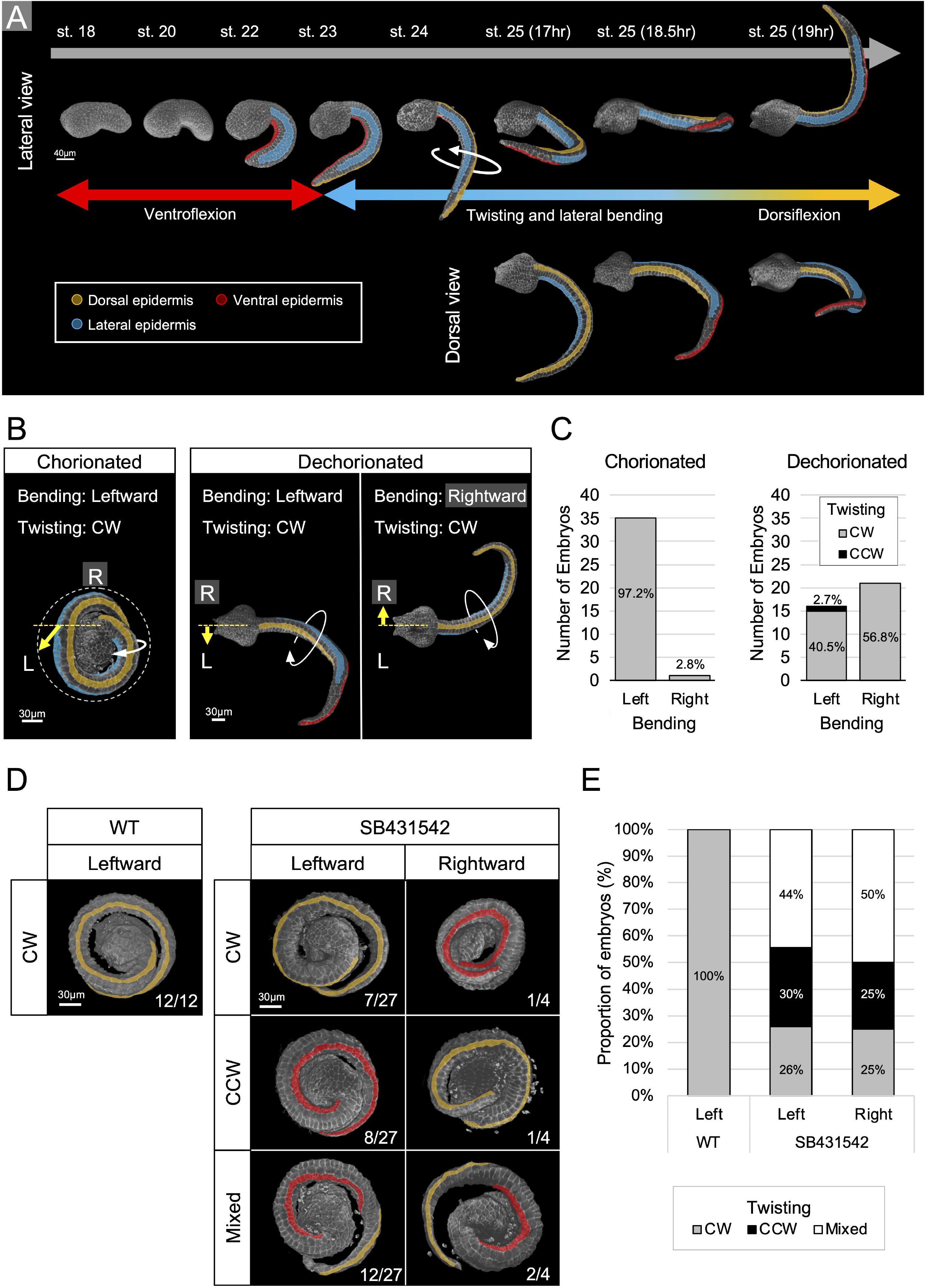
The axial rotation-like morphological change in *Ciona* consists of two components: bending and twisting. (A) Morphological changes during the tailbud stages in *Ciona*. Embryos were visualized by staining F-actin with phalloidin. The upper panels show lateral views, while the lower panels show dorsal views. The ventral midline epidermis, lateral epidermis, and dorsal midline epidermis are represented in red, light blue, and orange, respectively. Scale bar: 40 μm. (B, C) Bending and twisting directions in chorionated and dechorionated embryos. Embryos were fixed at st. 25 and stained with phalloidin. The bending direction was defined by the curvature of the anterior half of the tail relative to the trunk, as viewed from the adhesive papillae side. (B) The dotted line around the chorionated embryo indicates the region originally occupied by the chorion, which was manually removed. Yellow arrows indicate bending directions, while white arrows indicate twisting directions. The left panel shows a chorionated embryo in a left-sided view, and the right panel shows dechorionated embryos in a dorsal view. The ventral midline epidermis, lateral epidermis, and dorsal midline epidermis are shown in red, light blue, and orange, respectively. Scale bar: 30 μm. (C) The left and right graphs depict the bending direction. The gray and black bars indicate clockwise and counterclockwise twisting, respectively. CW: clockwise; CCW: counterclockwise. (D, E) Bending and twisting directions of WT and SB431542-treated embryos under chorionated conditions. Embryos were fixed at stage 25 and stained with phalloidin. (D) Embryos were oriented with the curved side of the tail (left or right relative to the trunk) facing forward. The ventral midline epidermis and dorsal midline epidermis are shown in red and orange, respectively. Scale bar: 30 μm. (E) Bar graph showing the proportions of each phenotype, normalized to 100% on the vertical axis. CW: clockwise; CCW: counterclockwise.

To further characterize these morphological changes, we examined the alignment of tail epidermal cell rows. The *Ciona* tail epidermis is organized into eight rows aligned along the anterior–posterior (AP) axis during the tailbud stage (Hotta et al., 2007; Kogure et al., 2022). This further observation revealed that the epidermis showed a twisted arrangement during the transitions in bending direction (Fig. 1A, st. 24–25; white arrow).

This progressive change in bending closely resembles AR observed in mice and rats, involving a 180° reversal of the bending axis along the lateral side, accompanied by axial twisting. Therefore, we refer to this phenomenon as AR in *Ciona* throughout this study.

### 3.2 Axial Rotation Consists of Twisting and Bending

Nodal, a member of the TGF-β superfamily, is a key factor in establishing left–right asymmetry in chordates (Hamada, 2020). Previous studies have also reported a correlation between Nodal signaling and AR in chordates (Collignon et al., 1996; Levin et al., 1997). In *Ciona*, Nodal is expressed on the left side of the lateral epidermis (Yoshida and Saiga, 2011, 2008). Dechorionation disrupts this pattern, resulting in bilateral Nodal expression (Yoshida and Saiga, 2008). Such dechorionation randomizes the bending direction at st. 23 (Palmquist and Davidson, 2017).

To determine whether dechorionation affects twisting, we analyzed embryos at st. 24–25, when twisting becomes evident. In chorionated embryos, the tail consistently bent to the left when viewed from the frontal side (Fig. 1B, C; n = 35/36). In dechorionated embryos, the tail bent randomly (Fig. 1B, C; leftward bending, n = 16/37; rightward bending, n = 21/37). In contrast, twisting remained clockwise in nearly all embryos, regardless of chorion status (Fig. 1B, C; chorionated, n = 36/36; dechorionated, n = 36/37). Since asymmetric Nodal expression is controlled by neurula rotation within the chorion (Nishide et al., 2012), we also observed embryos that were dechorionated after this event. As expected, these embryos exhibited leftward bending and clockwise twisting. These findings indicate that bending and twisting are regulated independently. The bending direction depends on the Nodal expression pattern (Nishide et al., 2012; Palmquist and Davidson, 2017), whereas twisting appears to be independent of this pattern.

We next investigated embryos treated with SB431542, an inhibitor of TGF-β signaling. Consistent with previous studies (Palmquist and Davidson, 2017), inhibition randomized the tail-bending direction in chorionated embryos (Fig. 1D, bending). Interestingly, TGF-β inhibition also disrupted the twisting direction. In leftward-bending embryos, twisting was clockwise (7/27), counterclockwise (8/27), or mixed, with both clockwise and counterclockwise twisting observed within the same embryo (12/27) (Fig. 1D, E; leftward-bending embryos treated with SB431542). In rightward-bending embryos, twisting was clockwise (1/4), counterclockwise (1/4), or mixed (2/4) (Fig. 1D, E; rightward-bending embryos treated with SB431542). These findings indicate that bending and twisting directions are uncorrelated under TGF-β signaling inhibition and that twisting direction depends on TGF-β signaling.

Together, these findings demonstrate that bending and twisting are two distinct processes. The bending direction is determined by Nodal expression (Nishide et al., 2012; Palmquist and Davidson, 2017), whereas the twisting direction is independently regulated through TGF-β signaling.

### 3.3 Quantification of the Twisting Angle During Axial Rotation

Although bending and twisting are distinct components, they occur simultaneously, making it difficult to quantify each separately. To evaluate twisting independently, we developed an image analysis method that mathematically extracts the twisting component from the curved 3D embryo shape (Fig. 2).

**Fig. 2.**
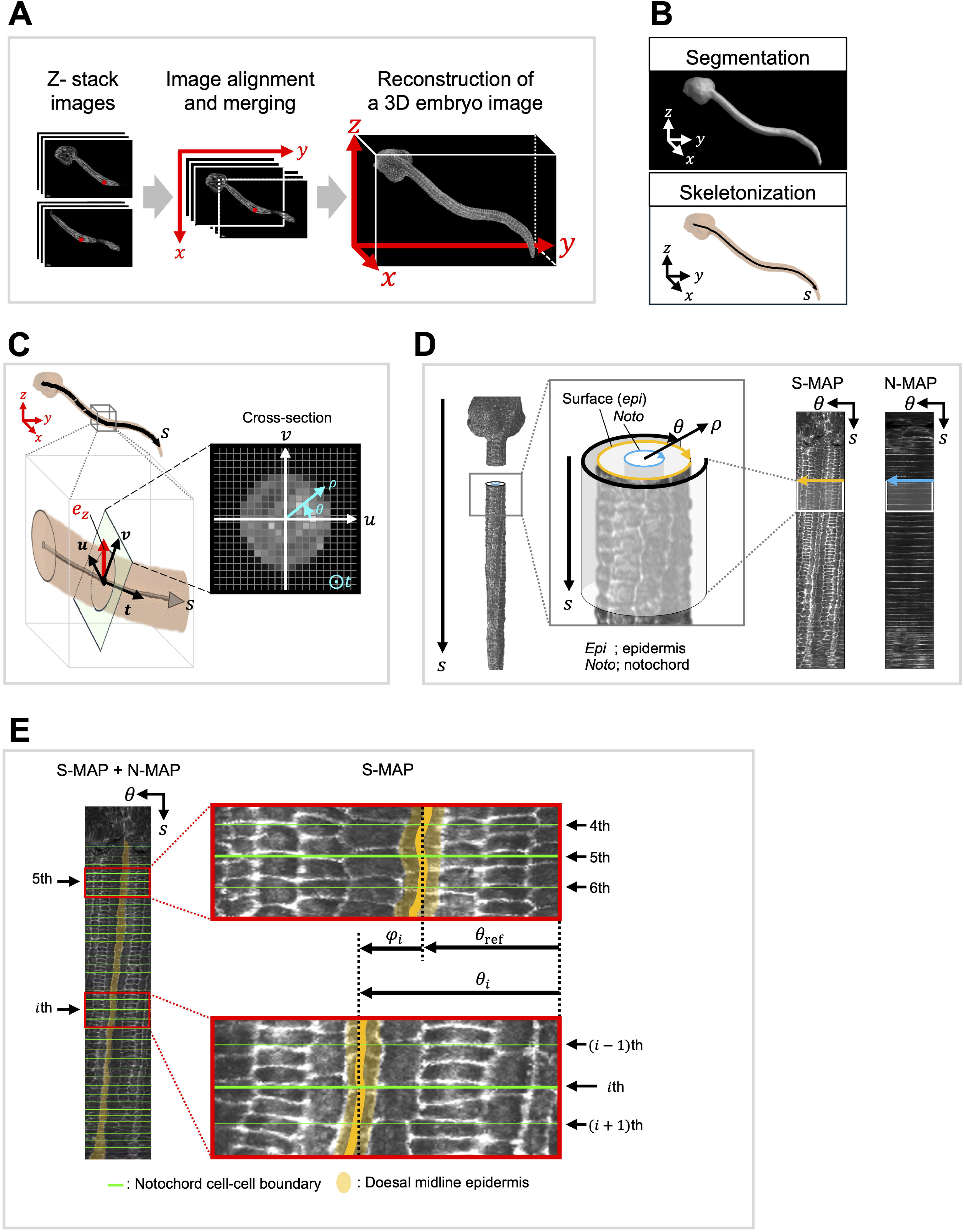
Schematic representation of the method used to quantify tail twisting. (A) Schematic of the overlap alignment procedure for large embryos that cannot fit within a single frame. Red dots indicate representative pixels in the overlapping regions. The *x*-, *y* -, and *z*-axes represent the spatial coordinates of the 3D images. (B) A 3D image of a segmented embryo and its skeleton. In the embryo image overlay (orange region), the black line marks the embryo’s skeleton, corresponding to its central axis . The *x* -, *y* -, and *z* -axes represent the spatial coordinates of the 3D image. (C) Schematic of cross-section determination. The *s*-axis represents the central axis. The vectors ***u, v***, and the tangent vector ***t*** are unit vectors that define the cross-sectional plane. The unit vector ***e***_*z*_ represents the *z*-axis of the spatial coordinate system, with ***v*** (*s*) defined to be as close as possible to the global *z*-axis, ***e***_*z*_ = (0,0,1). The *x*-, *y*-, and *z*-axes correspond to the spatial coordinates of the 3D images. The right panel shows a schematic of the cross-sectional image. The cross-sectional plane, defined by ***u, v***, and ***t*** in the local frame, is expressed in cylindrical-like coordinates (*ρ, θ,s*), where *ρ* is the radial distance from the centerline within the orthogonal plane, and *θ* is the angular direction in that plane. (D) Schematic of the procedure for generating S-MAP and N-MAP. The 3D image on the left depicts a virtual straight-tail embryo. The chain-like pattern shown in the S-MAP indicates the cell-cell boundaries of the epidermis. The ladder-like pattern shown in the N-MAP represents the notochord cell-cell boundaries. The *ρ*-, *θ*-, and *s*-axes correspond to a cylindrical coordinate system. (E) Schematic of the procedure for calculating the twisting angle. The green lines on the MAPs indicate the position of the notochord cell boundary. The dorsal midline epidermis is shown in orange region. The orange line in the right panel marks the midline of the dorsal midline epidermis. Each angular position *θ* within the neighboring regions, from the 4th to the 6th and (*i* − 1)-th to (*i* +1)-th, was used for smoothing to calculate *θ*_ref_ and *θ*_*i*_. Black arrows indicate the smoothed angular positions *θ*_ref_, *θ*_*i*_ and *φ*_*i*_ of each notochord cell–cell boundary.

Additionally, 3D reconstructions of entire embryos were generated within a 3D coordinate system (*x,y,z*)using Z-stack images of phalloidin-stained embryos fixed at several time points (Fig. 2A). For embryos exceeding the microscope’s field of view, images were acquired in partially overlapping segments and subsequently merged into a continuous Z-stack covering the entire embryo. To extract the anterior–posterior axis, the images were binarized to separate internal from external regions, followed by skeletonization that preserved topology, yielding a centerline *S* (Fig. 2B). At each position *s* along, the tangent vector ***t***(*s*) was defined, and cross-sections orthogonal to ***t***(*s*) were described by basis vectors ***u***(*s*) and ***v***(*s*) (Fig. 2C). To ensure consistent orientation, ***v***(*s*) was constrained to align closely with the *z*-axis of the *xyz* coordinate system (Fig. 2C). Using this local frame, the embryo was expressed in cylindrical-like coordinates (*ρ, θ,s*), where *ρ* denotes the radial distance from within the orthogonal plane, and *θ* represents the angular direction in that plane (Fig. 2C). Fluorescence intensity was resampled in this coordinate system, and the reoriented cross-sections produced a virtual straight-tail embryo, enabling quantification of the twisting component (Fig. 2D).

To visualize the intensity distributions on embryo surfaces, we generated S-MAPs (Fig. 2E). In the cylindrical coordinate system (*ρ, θ,s*), the embryo surface was identified as the outer boundary along the *ρ*-axis, and fluorescence intensity values at this boundary were projected onto a plane with (*θ,s*) coordinates. This procedure yielded S-MAPs in which curved surfaces were unwrapped into flat images, allowing direct comparison of epidermal rows along the AP axis. S-MAPs obtained after 14 hpf (at 16°C) revealed the arrangement of dorsal epidermal rows (Fig. 3A). The twisting angle was defined based on the positions of dorsal midline epidermal cells in the S-MAP.

**Fig. 3.**
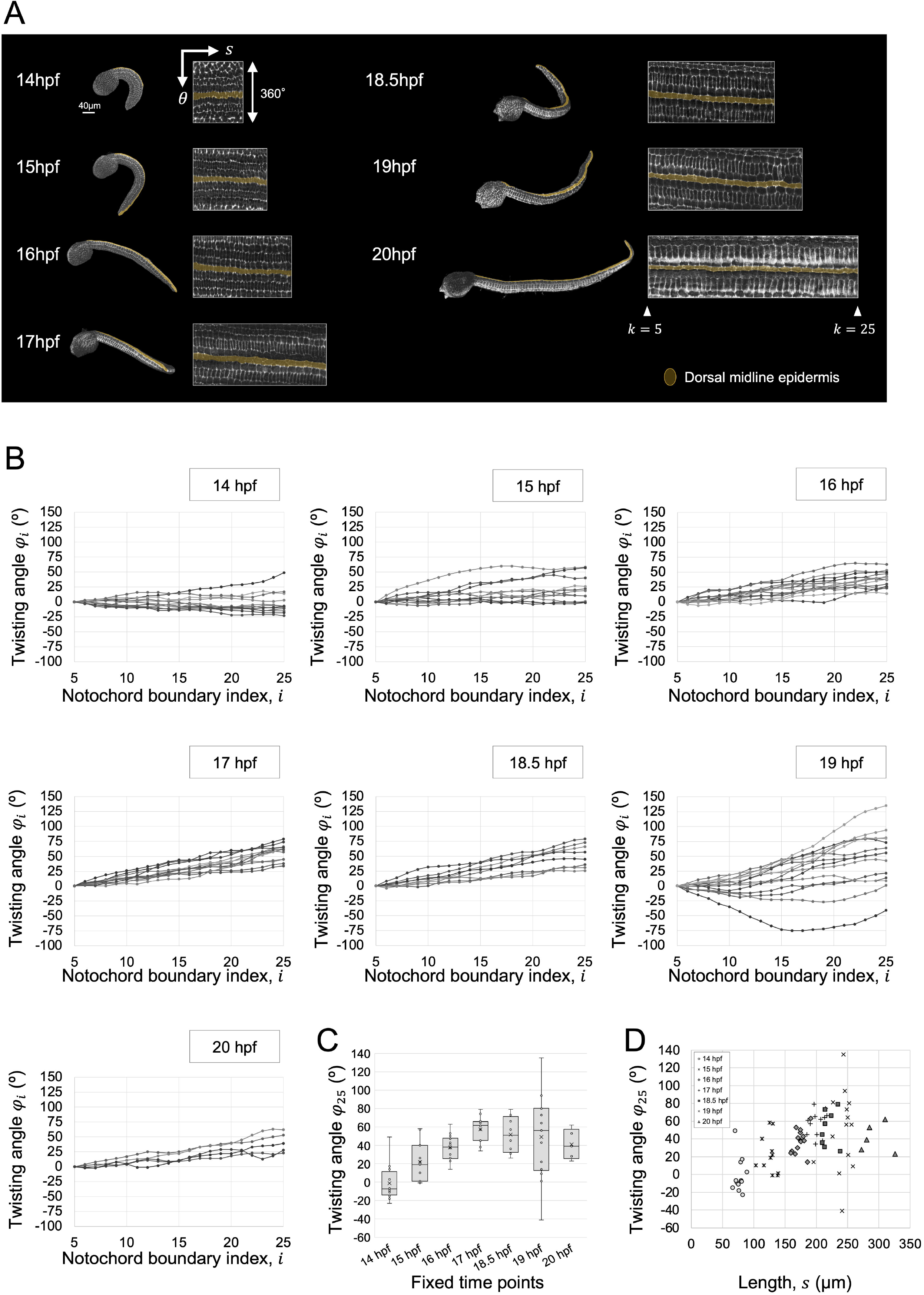
Twisting angle profiles across developmental stages. (A) The region from the 5th to the 25th notochord cell–cell boundaries in the S-MAPs, generated using the method described in Fig. 2, is shown alongside lateral views of their original 3D images. Each 3D image and map corresponds to embryos fixed at 14 hpf, 15 hpf, 16 hpf, 17 hpf, 18.5 hpf, 19 hpf, and 20 hpf, which developed at 16°C. The orange region indicates the dorsal midline epidermis. The left edge of the map represents the 5th boundary, while the right edge indicates the 25^th^ boundary . The *s* variables and *θ* describe the axes of the curvilinear coordinate system (*ρ, θ,s*)on the S-MAPs. (B) Twisting angle profiles at each notochord cell boundary index across developmental time points. Each graph represents a different developmental stage: 14 hpf (n = 12), 15 hpf (n = 11), 16 hpf (n = 14), 17 hpf (n = 12), 18.5 hpf (n = 8), 19 hpf (n = 14), and 20 hpf (n = 5). Line plots depict twisting angle profiles of individual embryos. The x-axis represents the notochord cell boundary index, and the *y*-axis indicates the twisting angle. Twisting angles were normalized by setting the angle at the 5th notochord cell boundary to 0°. Positive values indicate clockwise twisting. (C) Twisting angles at the 25th notochord cell-cell boundary at each developmental time point. Box plots display the twisting angle values at this boundary, derived from the data in (B). The box represents the interquartile range (IQR), with the horizontal line indicating the median. Whiskers extend to the minimum and maximum values within 1.5 × IQR. The cross (×) denotes the mean value. (D) Scatter plot of the length of the region from the 5th to the 25th notochord cell boundary versus the cumulative twisting angle within the same region. A strong positive correlation is observed between the twisting angle and region length from 14 hpf to 17 hpf (Spearman’s ρ = 0.77, p = 7.0 × 10?^11^ < 0.001).

Because notochord cells cease dividing after the mid-tailbud stage (Kugler et al., 2011), their boundaries provide stable landmarks. Therefore, we generated N-MAPs that display the positions of the boundaries between notochord cells as a ladder-like pattern along *S* (Fig. 2E). We then counted the number of notochord boundaries *i* from the anterior and defined the twisting angle at the *i* -th boundary as *φ*_*i*_ (Fig. 2E; green line). We also defined the twisting angle calculated every 10μm along *S* as *ϕ* _*j*_. Both *φ*_*i*_ and *ϕ*_*j*_ were referred to the 5th boundary, such that *φ*_5_ =0 and *ϕ* _0_=0 at this position.

At 14 hpf, dorsal midline epidermal rows were nearly parallel to the AP axis. However, as development progressed, these rows gradually slanted downward (Fig. 3A), suggesting progressive clockwise twisting. Correspondingly, twisting angle profiles (*φ*_*i*_, from *φ*_5_ to *φ*_25_) plotted against notochord cell–cell boundaries showed a linear relationship. This relationship was detectable in some embryos at 14–15 hpf but became robust and consistent in nearly all embryos from 16 hpf onward (Fig. 3B; linear regression, R^2^ > 0.7; 14 h: 6/12, 15 h: n = 7/11, 16 h: n = 13/14, 17 h: n = 12/12, 18.5 h: n = 8/8, 19 h: n = 9/14, 20 h: n = 5/5). Additionally, twisting angles calculated every 10 µm (*ϕ* _*j*_) along the tail AP axis in the same region (from the 5th to the 25th notochord boundary) also showed a linear relationship with tail length (data not shown; linear regression, R^2^ > 0.7; 14 h: 7/12, 15 h: n = 7/11, 16 h: n = 13/14, 17 h: n = 12/12, 18.5 h: n = 8/8, 19 h: n = 9/14, 20 h: n = 4/5). These data indicate that tail twisting is consistently distributed along the AP axis rather than being restricted to localized regions.

To examine developmental dynamics, we compared twisting angles at the 25th notochord boundary (*φ*_25_) across different stages. At 14 hpf, the average *φ*_25_ was close to zero (mean ± SD: –1° ±□20°), but it progressively increased in the clockwise direction, peaking at 17 hpf (58° ± 14°) before declining thereafter (Fig. 3C). At 19 hpf, unusually large fluctuations were observed compared to other stages (Fig. 3B-D); although the cause remains unclear, we note this as a characteristic feature. Finally, the axial length between the 5th and 25th boundaries showed a strong positive correlation with the twisting angle *φ*_25_ during 14–17 hpf (Fig. 3D; Spearman’s ρ = 0.76, p < 0.001).

### 3.4 Tail Twists Occur Together with Internal Tissue Movements, Independent of the Trunk or Tail Tip

While S-MAP allowed quantification of the twisting angle in the tail epidermis, it remained unclear whether this deformation was limited to the surface or extended throughout the entire tail. The tail consists of internal tissues—including muscle, endoderm, and neural tube—that surround the notochord and are covered by epidermal cells (Fig. 4A). To test whether internal tissues also undergo twisting, we examined cross-sectional images of the tail at the 5th, 15th, and 25th notochord cell–cell boundaries. The relative positions of the epidermis and internal tissues were preserved in all three cross-sectional images (Fig. 4B, n = 13), indicating that not only the epidermis but also the muscle, endoderm, and neural tube twist along the AP axis.

**Fig. 4.**
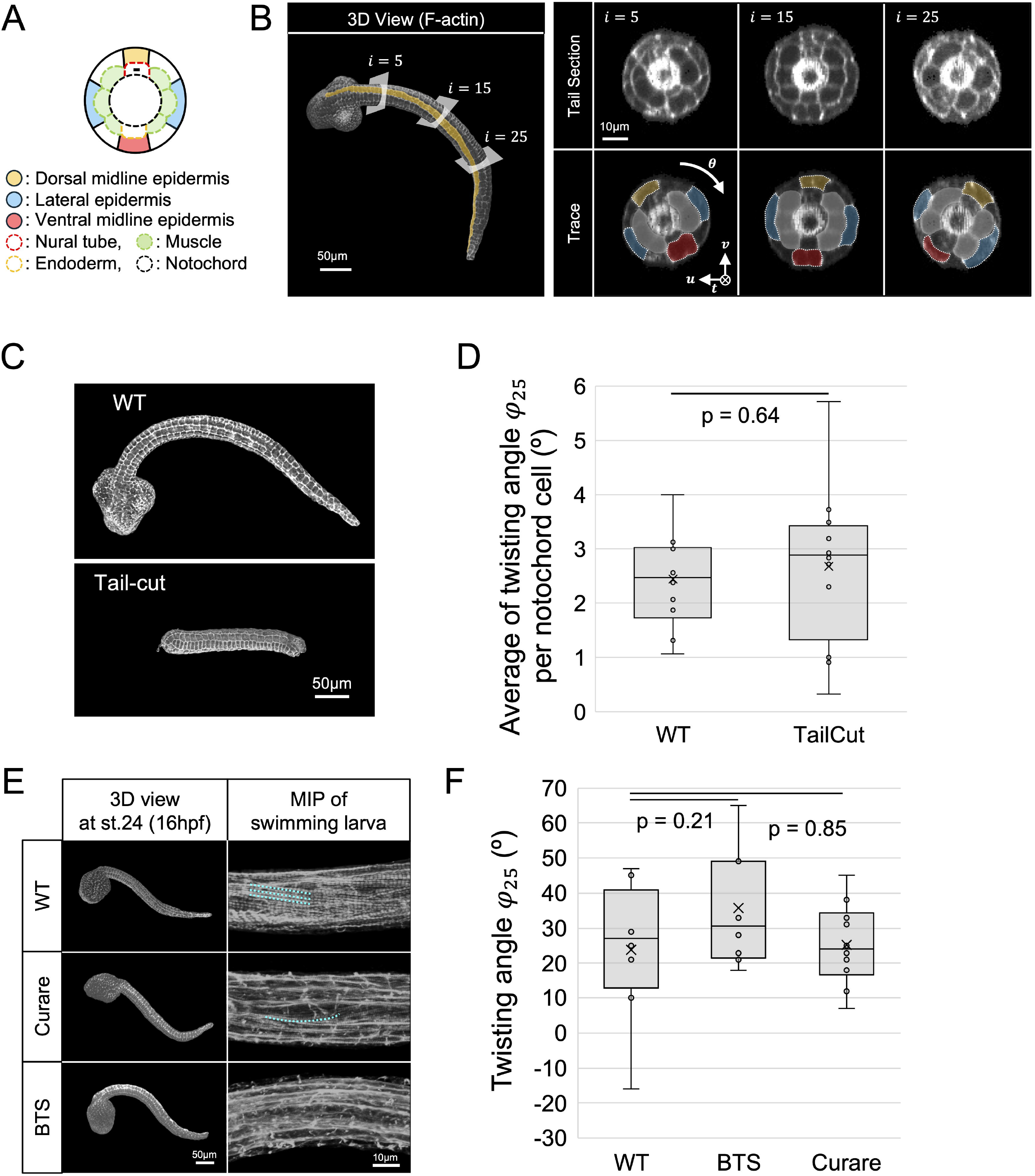
Tail twisting emerges independently within the tail domain. (A) Schematic illustration of a cross-section of the tail region. (B) A tailbud-stage embryo fixed at 17 hpf was stained with phalloidin. The embryo was developed at 16°C. Cross-sections were obtained at the boundaries of the 5th, 15th, and 25th notochord cells. Red indicates ventral midline epidermal cells, blue indicates lateral epidermal cells, orange indicates dorsal midline epidermal cells, and white indicates muscle cells. A, anterior; P, posterior; D, dorsal; V, ventral; R, right; L, left. Scale bar: 50 μm for the 3D view panels; 10 μm for the tail section and trace panels. (C) A WT tailbud-stage embryo and a tail fragment (tail-cut), both fixed at 16 hpf, were stained with phalloidin. Embryos were incubated at 16°C. Scale bar: 50 μm. (D) Box plots illustrating the distribution of average twisting angles per notochord cell in WT and tail-cut embryos. For WT embryos, the average was calculated using the 11th to 25th notochord cells, corresponding to the region retained in the tail-cut condition. The box represents the interquartile range (IQR), with the horizontal line indicating the median. Whiskers denote the minimum and maximum values within 1.5 × IQR. The cross (×) indicates the mean value. Statistical comparisons were conducted using Welch’s t-test; no significant difference was detected (Welch’s test, p = 0.64 > 0.05). (E) Embryos fixed at 16 hpf or at the swimming larva stage were stained with phalloidin. All embryos were incubated at 16°C. The left panels present a 3D view at 16 hpf, while the right panels display maximum intensity projections (MIPs) of swimming larvae. The dotted line indicates the myofibril pattern. Scale bars: left panels, 50 μm; right panels, 10 μm. (F) Box plots depicting the distribution of twisting angles in WT, BTS-treated, and curare-treated embryos. The accumulated angle was calculated from the 5th to the 25th notochord boundary. The box represents the interquartile range (IQR), with the horizontal line indicating the median. Whiskers extend to the minimum and maximum values within 1.5 × IQR. The cross (×) denotes the mean value. Statistical comparisons were performed using Welch’s t-test; no significant differences were observed between WT and BTS (Welch’s test, p = 0.21 > 0.05) or between WT and curare (Welch’s test, p = 0.85 > 0.05).

To determine whether tail twisting occurs independently within the tail region, we surgically removed the trunk and tail tip at 13 hpf, prior to the onset of visible twisting (Fig. 4C). For analysis, we used embryos fixed at 17 hpf that had eight epidermal cell rows. To avoid potential artifacts caused by the physical impact of cutting, we excluded several notochord cells adjacent to the cut site from the analysis. Accordingly, in WT embryos, the region from the 10th to the 25th notochord cells was analyzed. Clockwise twisting was observed in both WT and tail-cut embryos. The average twisting angle, computed per notochord cell, showed no significant difference between WT (n = 10) and tail-cut (n = 12) embryos (Fig. 4D). These results indicate that clockwise twisting can occur independently within the isolated tail fragment.

Previous studies have reported that tail muscle cells, a component of tail tissue, acquire left–right asymmetric alignment of myofibrils during development (Matsuo et al., 2021). To test whether this alignment contributes to twisting, we treated embryos with curare (d-tubocurarine chloride), an acetylcholine receptor blocker, and BTS, an actomyosin ATPase inhibitor. Consistent with previous findings (Ohtsuka and Okamura, 2007), WT embryos showed organized myofibril patterning (n = 6), whereas both treatments disrupted myofibril patterning at the swimming larval stage (Fig. 4E; curare, n = 7; BTS, n = 8). However, neither the twisting direction nor the twisting angle (*φ*_25_) at 16 hpf differed significantly between treated and wild-type embryos (Fig. 4F; WT, n = 8; BTS, n = 8; curare, n = 10). These findings indicate that left-right asymmetric myofibril patterning is not essential for either the generation or directional control of twisting. Thus, tail twisting represents an intrinsic property of the tail as a whole, occurring together with internal tissues and independently of trunk interactions or myofibril asymmetry.

### 3.5 TGF-β Signaling Coordinates Local Twisting Directions Along the Tail

To further investigate the role of TGF-β signaling in twisting, we analyzed SB431542-treated embryos using our method which quantifies twisting angles (Fig. 2). Treated embryos exhibited shortened tails, incomplete notochord intercalation, and a swollen mid-tail resembling a “tsuchinoko,” a snake-like creature from Japanese folklore (Fig. 5A). Because notochord intercalation failed, the notochord cell–cell boundaries could not serve as reliable landmarks. Therefore, we measured twisting angles *ϕ* along a 150 μm AP region of the tail, calculating the angle at every 10 µm interval.

**Fig. 5.**
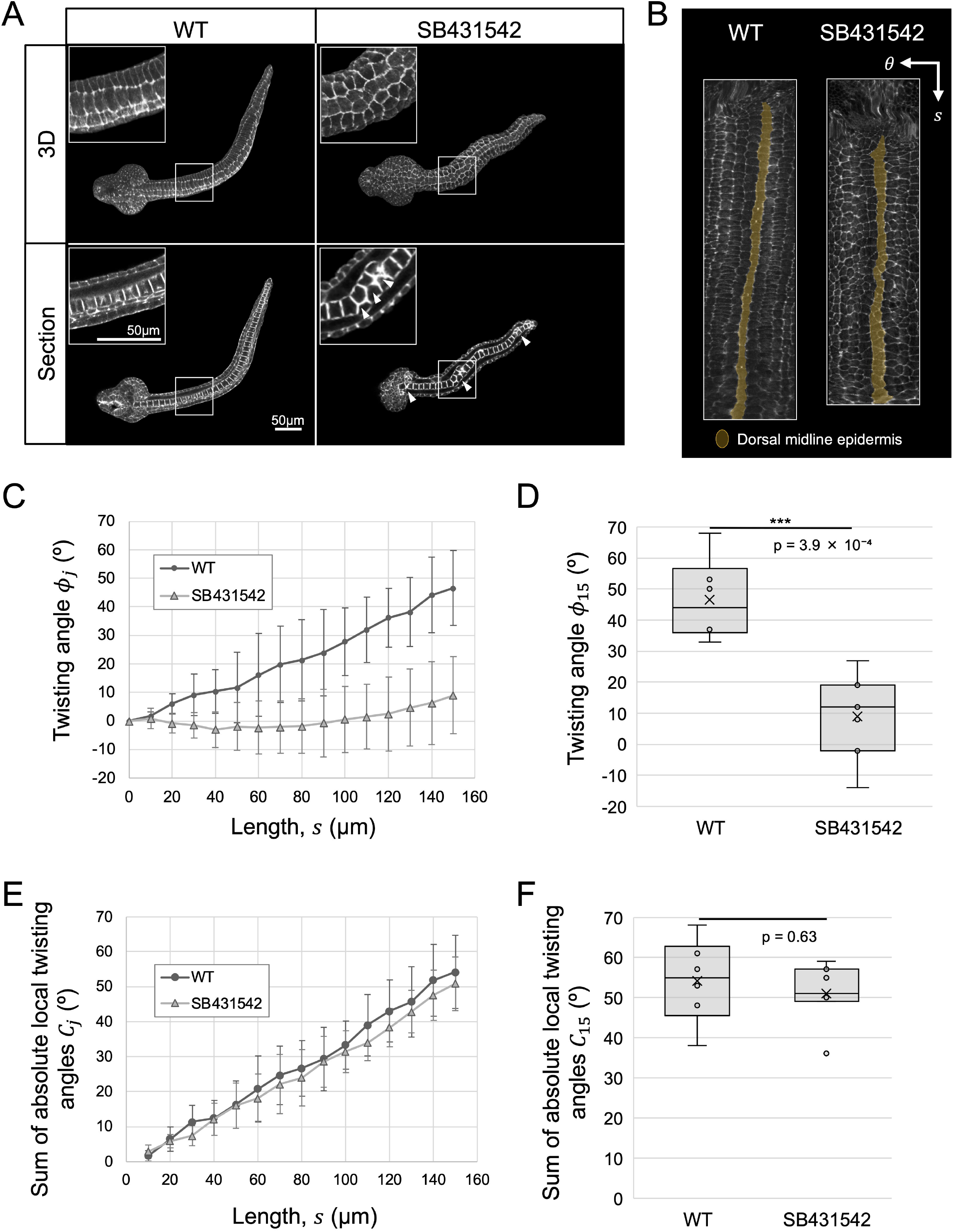
TGF-β signaling unifies the direction of twisting. (A) Embryos were incubated at 16°C, fixed at 16 hpf, and stained with phalloidin. SB431542 was added at the late neurula stage at a final concentration of 2.5 µM and maintained in the medium until just before fixation. The upper panels display 3D reconstructions, while the lower panels show sectional images. In the sectional images, notochord cells appear as ladder-like structures. In SB431542-treated embryos, notochord cell intercalation appears incomplete (arrowheads). Scale bar: 50 μm. (B) 2D surface projection maps of WT and SB431542-treated embryos. Orange regions indicate the dorsal midline epidermis. *θ* and *s* represent the curvilinear coordinate system (*ρ, θ,s*)on S-MAPs. (C) Twisting angles of WT (n = 6) and SB431542-treated embryos (n = 7) measured from the corresponding 5th notochord boundary. The *Y*-axis represents the twisting angle, and the *x*-axis indicates the length of the region. Error bars denote the standard deviation (SD). (D) Box plot showing the twisting angles at 150 µm, extracted from the data in (C). The box represents the interquartile range (IQR), with the horizontal line indicating the median. Whiskers extend to the minimum and maximum values within 1.5 × IQR. The cross (×) denotes the mean value. A significant difference was detected between WT and SB431542-treated embryos (Welch’s t-test, p = 3.9 × 10?L < 0.001). (E) The sum of absolute twisting-angle differences calculated from data shown in (C). Angles were calculated for every 10 µm segment along the *s*-axis. Error bars represent the standard deviation (SD). (F) Box plot showing the sum of absolute twisting-angle differences at 150 µm, extracted from the data in (E). The box represents the interquartile range (IQR), with the horizontal line indicating the median. Whiskers extend to the minimum and maximum values within 1.5 × IQR. The cross (×) denotes the mean value. No significant difference was detected between WT and SB431542-treated embryos (Welch’s t-test, p = 0.63 > 0.05).

Inhibition of TGF-β signaling disorganized the local twisting direction, resulting in a reduced twisting angle *ϕ* across the analyzed region (Fig. 5B-D). Nevertheless, twisting itself was still observed. To determine whether inhibition affected both the direction and magnitude of twisting, we defined the absolute local twisting angle (|Δ *ϕ*_*j*_ |) as the absolute angular difference between consecutive 10-µm intervals, and summed these values along the tail starting from the *s* corresponding to the 5th notochord position (*C*_*j*_). Interestingly, there was no significant difference in the total twisting magnitude over the 150 µm AP region of the tail between WT and SB431542-treated embryos (Fig. 5E, F), indicating that inhibition randomized the twisting directions but did not reduce the magnitude of local twisting. These results suggest that TGF-β signaling may be not required to generate twisting per se but is necessary to align local twisting into a coordinated global direction along the AP axis.

## 4. Discussion

A schematic summary of AR in *Ciona* is presented in Fig. 6. This study demonstrates that AR consists of two components. The first component, bending, involves a sequential shift in tail posture—from ventral to lateral to dorsal—culminating in a 180° rotation of the tail tip around the trunk (upper part of Fig. 6). Previous work suggests that this bending direction depends on Nodal expression (Nishide et al., 2012; Palmquist and Davidson, 2017). The second component, twisting, is described as a clockwise twisting of the tail along the AP axis. We established a method to quantify twisting independently of bending. Using this method, we found that the average twisting angle increased to approximately 60° by 17 hpf and then declined (Fig. 3B). Notably, the twisting angle plotted against notochord cell–cell boundaries or length along the AP axis showed linear relationship, indicating that twisting progresses uniformly along the measured region rather than being localized. We also revealed that TGF-β signaling is necessary for coordinating the directionality of this uniform twisting.

**Fig. 6.**
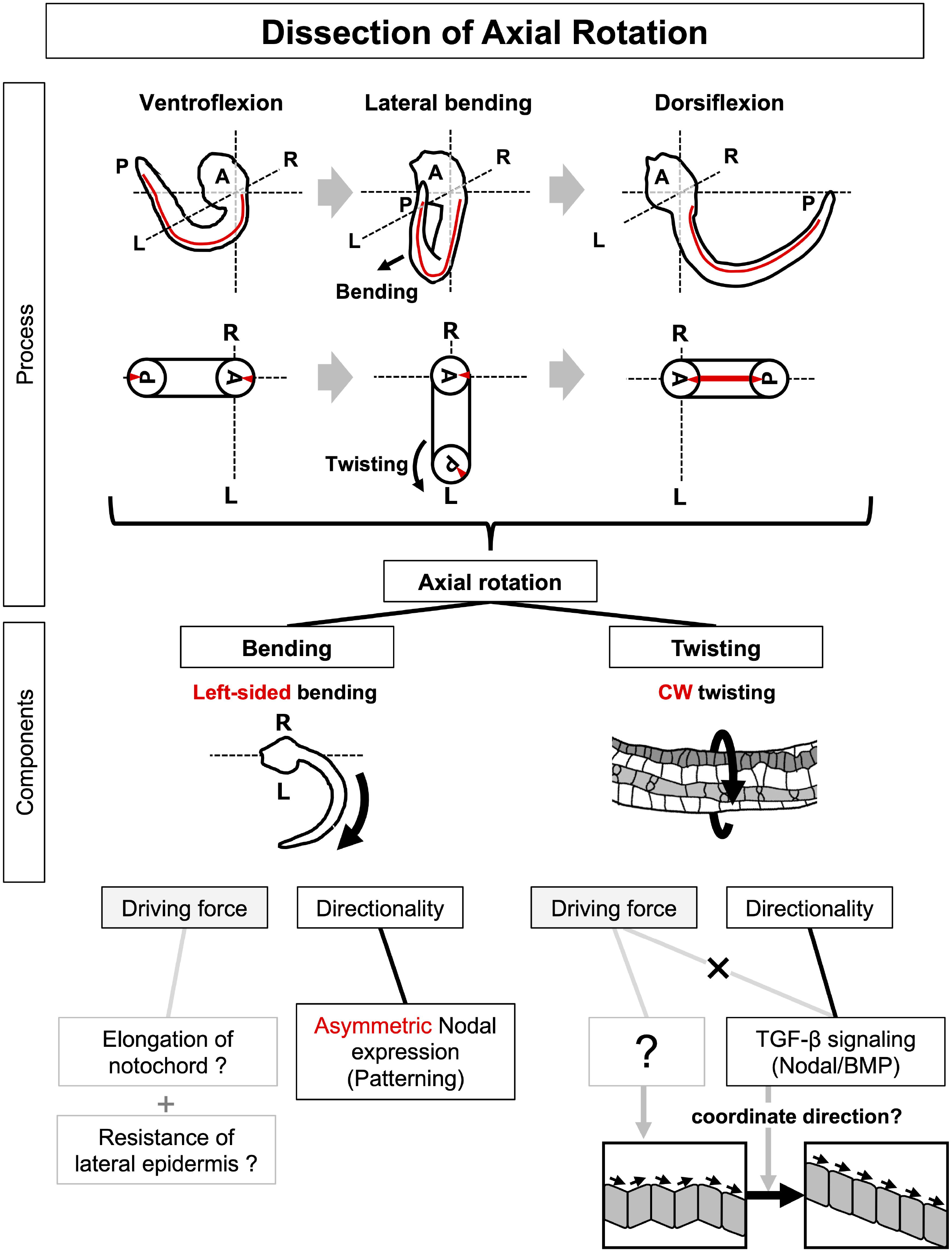
Proposed model illustrating the regulation of axial rotation in the *Ciona* tailed embryo. This schematic model summarizes AR in Ciona. We have dissected AR into two components: twisting and bending. The upper schematic images illustrate the embryo’s morphological transition during AR, showing the tail bending from ventral to lateral to dorsal positions while twisting. Each component can be further subdivided into driving force and directionality. Both the directionality of bending and twisting are regulated by TGF-β signaling, which may coordinate local twisting directions into a coherent global pattern.

### 4.1 Regulation of Bending During AR

Previous studies have shown that bending laterality depends on Nodal expression (Nishide et al., 2012; Palmquist and Davidson, 2017). We confirmed that disrupting asymmetric Nodal signaling—either by dechorionation or TGF-β inhibition—randomizes the bending direction (Fig. 2). While prior studies emphasized specific developmental stages, our analysis revealed a continuous shift in bending direction: ventral → lateral → dorsal (Fig. 1A; Suppl. Fig. 2A).

Ventral bending in *Ciona* has been attributed to resistance of ventral epidermal cells to notochord elongation, a process regulated by Admp/Bmp signaling(Kogure et al., 2022). BMP signaling is active in the ventral epidermis from the late gastrula to early tailbud stages (Kogure et al., 2022; Waki et al., 2015). In contrast, lateral bending depends on left-sided Nodal expression, which arises later during the tailbud stage (Imai et al., 2004; Yoshida and Saiga, 2008); https://aniseed.fr/). This temporal sequence—BMP activity followed by Nodal expression—may underlie the transition from ventral to lateral bending (Fig. 2). However, the molecular mechanism driving the final dorsal bending remains unknown.

Downstream of Nodal, Pitx2 is also expressed in the left lateral epidermis during the mid-tailbud stage (Yoshida and Saiga, 2008). A recent study on chick gut morphogenesis showed that Pitx2 increases tissue stiffness on the expressing side, contributing to asymmetric gut tilting (Sanketi et al., 2022). By analogy, lateral bending in *Ciona* may result from the resistance of the lateral epidermis to notochord elongation, regulated by asymmetric Nodal–Pitx2 expression.

The importance of the notochord has also been suggested in mouse AR. Mouse mutants that fail to undergo AR, such as the *not turning* mutants (Melloy et al., 1998) and *rotatinGT* mutants (Faisst et al., 2002), display notochordal defects. This may explain why embryos with bilateral Nodal expression or inhibited Nodal signaling show randomized lateral bending (Fig. 2). Since bending is highly sensitive to mechanical asymmetries, even subtle differences in epidermal stiffness could influence the direction of bending. Future measurements of epidermal mechanical properties may help test this hypothesis.

### 4.2 Regulation of Twisting During AR

In contrast to the bending component, the tail consistently twisted clockwise, even in dechorionated embryos (Fig. 1B, C). This observation suggests that twisting occurs independently of the asymmetric Nodal expression pattern. Interestingly, inhibition of TGF-β signaling disrupted the global consistency of twisting (Fig. 5). Since the sum of the absolute twisting magnitudes did not differ significantly from WT (Fig. 5B, C), we propose that TGF-β signaling is not required to generate the twisting force. Instead, it is necessary to align local twisting into a coordinated global direction along the AP axis (Fig. 5, 6). TGF-β signaling and its downstream effectors are known regulators of cell polarity during morphogenetic movements and cell shape changes. For example, in the zebrafish heart tube, Nodal promotes rotation by directing cell rearrangements (Kidokoro et al., 2022). BMP signaling regulates polarized cell shape formation in the *Ciona* ventral epidermis (Kogure et al., 2022). Pitx2, a Nodal target, converts mesendodermal cell movements into directional migration during zebrafish gastrulation (Collins et al., 2018). These examples support the idea that directional coordination by TGF-β signaling is functionally and evolutionarily conserved. Further investigation of downstream effectors, using our analysis method to quantify twisting independently of bending, may provide deeper insights into the molecular and mechanical mechanisms that coordinate asymmetry and enable a trans-scale understanding of AR from single cells to the organismal level.

Cross-sectional analyses and tail-cut experiments (Fig. 4B-D) showed that clockwise twisting can occur independently within isolated tail fragments. This suggests that the epidermis, muscles, neural tube, endodermal strand, and notochord are candidate tissues involved in generating twisting. The strong linearity of twisting angles along the AP axis (Fig. 3A) further implies regulation at the level of small tissue units. In AR of rat and chicken embryos, twisting-like events occur in temporally and regionally restricted patterns, initiating in the upper body and progressing caudally (Fujinaga et al., 1995; Hamburger and Hamilton, 1951). This difference may reflect that our analysis focused on the limited tail region of *Ciona*, so temporal or regional changes elsewhere may not have been detected. Future 4D imaging of the entire embryo will be necessary to determine whether the spatial progression of twisting also occurs in *Ciona*.

Previous studies in mice have linked AR to notochord formation (Faisst et al., 2002), dorsoventral somite asymmetry (Matsuda, 1991) and asymmetric neural tube cell division (Poelmann et al., 1987). In *Ciona*, TGF-β–inhibited embryos displayed incomplete notochord formation yet still underwent twisting, suggesting that notochord formation is not essential for generating the twisting force. Muscles, which are bilaterally positioned in the tail (Nakamura et al., 2012) and drive lateral tail movements (McHenry, 2005) have also been considered potential contributors. Left–right asymmetric myofibril twisting in the tail muscle has been reported (Matsuo et al., 2021), raising the possibility of muscle involvement in AR. However, when myofibril patterning was disrupted, twisting persisted at wild-type levels (Fig. 4E, F), indicating that myofibril patterning is not essential for generating or directing twisting. While several tissues may contribute, the precise driver remains unknown. Future tissue-level manipulations will help identify the source of the twisting force during AR.

### 4.3 Evolutionary Conservation of Axial Rotation in Chordates

In *Ciona*, we showed that TGF-β signaling regulates both twisting and bending. The TGF-β family member Nodal has also been implicated in AR laterality in mouse and chick embryos (Collignon et al., 1996; Levin et al., 1997), suggesting evolutionary conservation among chordates. Building on our identification of bending and twisting as distinct components in *Ciona*, we should re-examine AR descriptions in other species.

In mouse and rat embryos, AR begins with dorsiflexion, followed by the tail shifting along the right side of the trunk and transitioning to ventroflexion, resulting in a counterclockwise 180° body axis rotation (Suppl. Fig. 1; Fujinaga et al., 1995; Poelmann et al., 1987). In mice, AR correlates with asymmetric Nodal expression (Collignon et al., 1996), similar to the bending observed in *Ciona*. However, *Ciona* bends leftward, while mice bend rightward. This opposite direction may reflect species-specific regulatory mechanisms linked to the spatial domains of Nodal expression: the left epidermis in *Ciona* (Yoshida and Saiga, 2008) versus the lateral plate mesoderm in mice (Shiratori and Hamada, 2006). Twisting has been repoted in rats, where the upper, middle, and lower body regions twist sequentially (Fujinaga et al., 1995). These findings suggest that AR in rodents may involve both bending and twisting.

In chicken embryos, AR has been associated with Pitx2, a Nodal downstream effector (Logan et al., 1998), suggesting that TGF-β signaling involvement in AR may be conserved between chickens and ascidians. According to the staging of (Hamburger and Hamilton, 1951), chicken AR begins with leftward head turning that progresses caudally until the tail completes its rotation. Unlike in mice, however, clear left–right bending is not observed; instead, the body axis appears to twist, indicating that twisting may predominate in chicken AR. Classical embryonic illustrations (Hamburger and Hamilton, 1951) hint that slight curvature may precede head turning, but whether both bending and twisting occur remains unclear.

It remains unresolved whether bending and twisting are independently regulated across species. Applying these concepts, coupled with our quantitative analysis, may provide new insights into the conserved and divergent mechanisms of AR.

### 4.4 The Role of AR in *Ciona*

AR may have evolved to help embryos adjust their posture within physically constrained environments.

In *Ciona*, AR may facilitate hatching by reorienting the embryo’s posture to enhance the effectiveness of tail flick movements. Although the link between AR and tail flicking remains unclear, flicking occurs before hatching (Akahoshi et al., 2021) and embryos position their lateral side against the chorion prior to hatching (Fig. 2D, WT; Suppl. Mov. 1). This posture may allow flicking to exert greater force against the chorion. These observations suggest that AR contributes to chorion rupture by establishing a favorable orientation for flick-driven mechanical interactions. Further research is needed to determine whether posture affects hatching success and whether tail flicking is essential for this process.

### 4.5 *Ciona* as a Valuable Model for Understanding AR

Recent efforts to generate synthetic mouse embryos have shown that development halts at embryonic day 8.5 (E8.5), corresponding to AR onset *in vivo* (Tarazi et al., 2022). This finding suggests that understanding AR mechanisms may be key to overcoming this developmental bottleneck.

Studying AR in vertebrate models is complicated due to lengthy developmental timelines and complex tissue structures. In contrast, *Ciona* embryos offer major advantages: they develop rapidly, possess a simple body plan, and consist of relatively few cells, enabling single-cell–level observation and quantification of morphogenetic changes across the entire embryo (Guignard et al., 2020; Hotta et al., 2007; Nakamura et al., 2012). Moreover, because *Ciona* embryos can develop without a chorion, they are particularly well suited for live imaging. These technical advantages make ascidians an ideal model for dissecting AR.

Taken together, our findings establish ascidians as a tractable and informative system for investigating AR and lay the groundwork for future studies aimed at uncovering its mechanistic basis.

## Supporting information

Suppl. Mov. 1

## Acknowledgments

*Ciona intestinalis* adults were provided by Dr. Yutaka Satou (Kyoto University), Dr. Manabu Yoshida (The University of Tokyo), and Dr. Yasunori Sasakura (Tsukuba University), with support from the National Bio-Resource Project of AMED, Japan.

## Competing Interests

The authors declare no competing financial interests.

## Author Contributions

Y.S.K. observed AR in *Ciona*. Y.S.K. and K.H. designed and performed the research. Y.S.K. and S.O. developed the MATLAB program, and Y.S.K. analyzed the data. Y.S.K. and K.H. wrote the manuscript, which K.O. critically revised. K.H. supervised the research.

## Funding

This work was supported by JSPS KAKENHI grants 23H04717, 24K02038, and 25H01799 awarded to K.H., as well as by JSPS KAKENHI grant 24KJ1934, SPRING JPMJSP2123, and the Keio University Doctorate Student Grant-in-Aid Program from the Ushioda Memorial Fund awarded to Y.S.K.

## Data Availability

All data generated or analyzed during this study are included in the manuscript and its supporting files.

## Figure Legend

**Suppl. Fig. 1. Schematic representation of axial rotation in the mouse embryo** (Gavrilov and Lacy, 2013).

Mouse embryos undergo dorsiflexion, followed by tail shifting along the right side of the trunk, transitioning into ventroflexion. This sequence results in a counterclockwise 180° body-axis rotation. A: anterior; P: posterior; red line: dorsal neural tube.

**Suppl. Fig. 2. Observation of tail bending direction**.

(A) Tail bending direction at multiple time points. From a frontal view of the cross-section at the 5th notochord boundary, the position of the tail tip relative to the center of this boundary was measured to define the bending direction, with the ventral direction set at 0°. The bending direction shifted from ventral to lateral and then to dorsal between 16 and 19 hpf. D: dorsal; V: ventral; A: anterior; P: posterior; L: left; R: right.

(B) Bending and twisting directions of embryos in which Nodal was expressed on the left side. Embryos were dechorionated after the establishment of left-sided Nodal expression (Nishide et al., 2012), from the early to late tailbud stage. All embryos (n = 11) exhibited leftward bending and clockwise twisting.

**Suppl. Mov. 1. Tail flicking**.

The embryo exhibited tail flicking in a radial direction (n = 1). It was cultured with the chorion until just before observation at 16 hpf and was manually dechorionated prior to imaging. The timestamp indicates the elapsed time after fertilization.

## Notes

### Competing Interest Statement

The authors have declared no competing interest.

