## Supplementary figures and images for "Axial Rotation Comprises Concurrent Twisting and Bending, Each Differentially Regulated by TGF-β Signaling"

### Suppl. Fig. 1

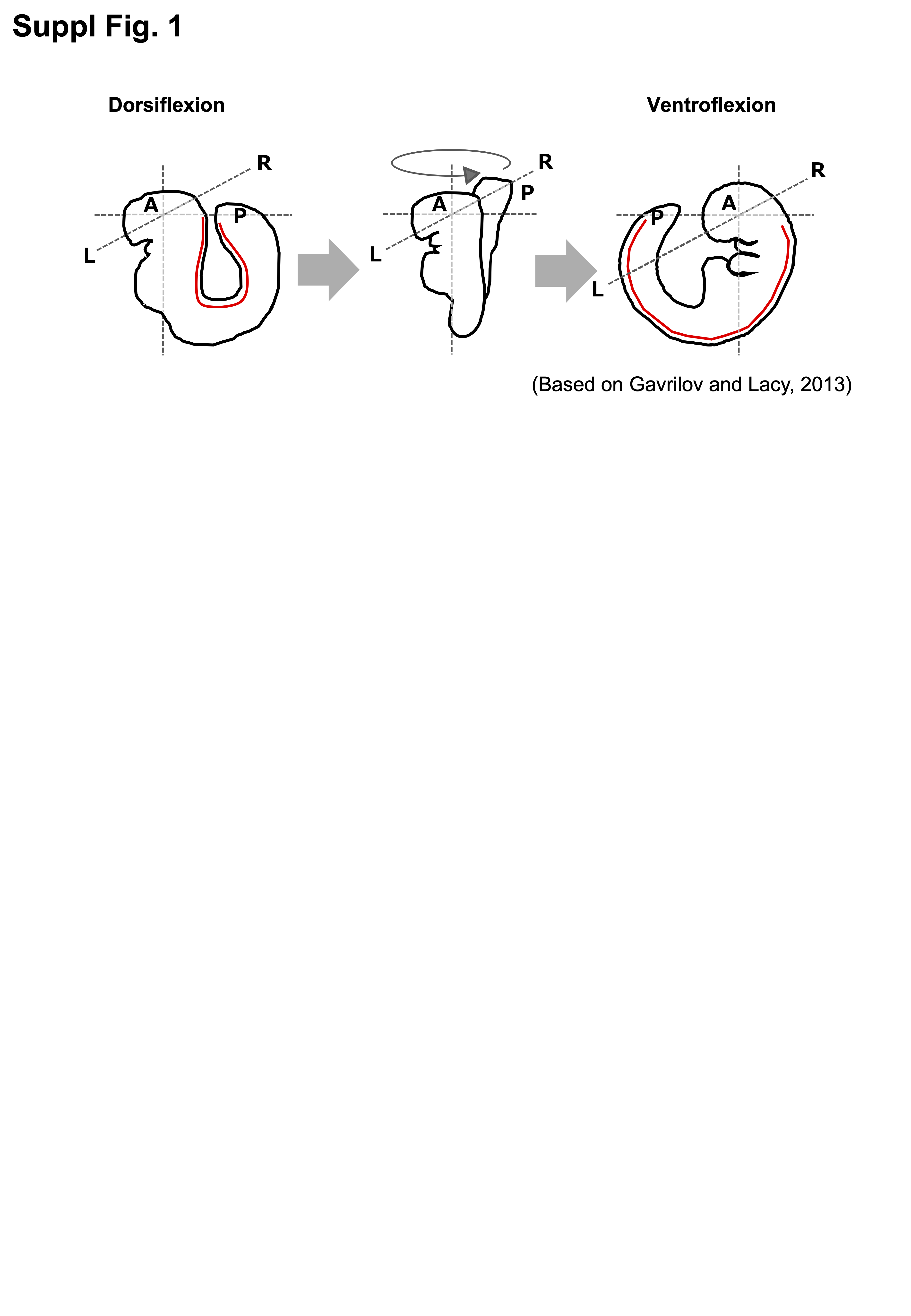

### Suppl. Fig. 2

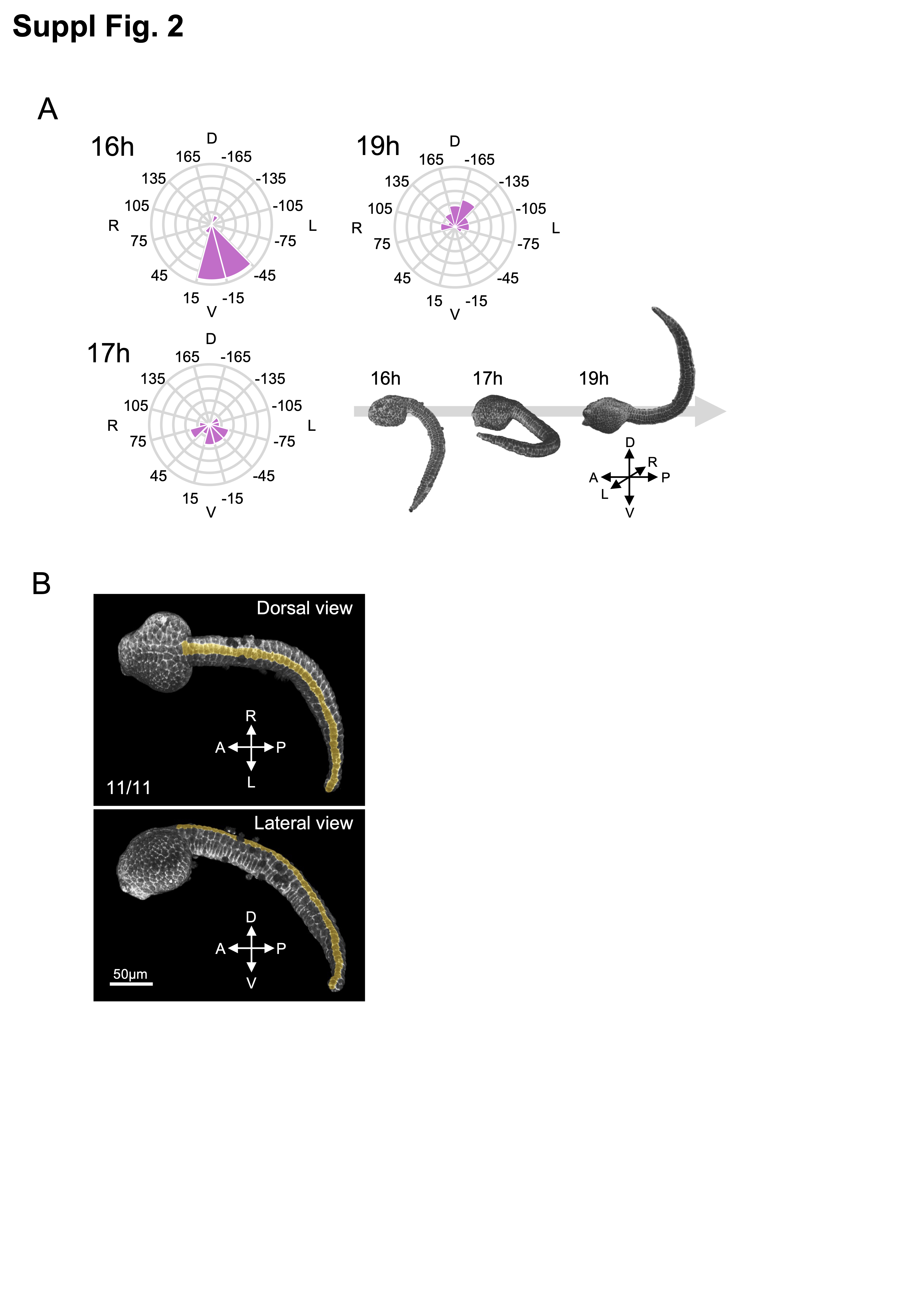
